# Functional connectivity differences in early infancy precede autism symptoms: a multivariate pattern analysis

**DOI:** 10.1101/866939

**Authors:** Abigail Dickinson, Manjari Daniel, Andrew Marin, Bilwaj Goanker, Mirella Dapretto, Nicole M. McDonald, Shafali Jeste

## Abstract

Functional brain connectivity is altered in children and adults with autism spectrum disorder (ASD). Mapping pre-symptomatic functional disruptions in ASD could identify infants based on neural risk, providing a crucial opportunity to mediate outcomes before behavioral symptoms emerge.

Here we quantify functional connectivity using scalable EEG measures of oscillatory phase coherence (6-12Hz). Infants at high and low familial risk for ASD (N=65) underwent an EEG recording at 3 months of age and were assessed for ASD symptoms at 18 months using the Autism Diagnostic Observation Schedule-Toddler Module. Multivariate pattern analysis was used to examine early functional patterns that are associated with later ASD symptoms.

Support vector regression (SVR) algorithms accurately predicted observed ASD symptoms at 18 months from EEG data at 3 months (*r*=0.76, *p*=0.02). Specifically, lower frontal connectivity and higher right temporo-parietal connectivity predicted higher ASD symptoms. The SVR model did not predict non-verbal cognitive abilities at 18 months (*r*=0.15, *p*=0.36), suggesting specificity of these brain alterations to ASD.

These data suggest that frontal and temporo-parietal dysconnectivity play important roles in the early pathophysiology of ASD. Early functional differences in ASD can be captured using EEG during infancy and may inform much-needed advancements in the early detection of ASD.

## Introduction

Autism is a disorder of early brain development that is diagnosed based on the presence of social-communication impairments and restricted, repetitive behaviors [1]. Interventions that begin early in life hold immense potential for altering neurodevelopmental trajectories and improving outcomes in autism spectrum disorder (ASD). However, the behavioral signs of ASD are typically identified after four years of age [2, 3], therefore preventing earlier attempts to mediate outcomes. Mapping the changes in early brain development that lead to ASD could inform pre-symptomatic markers of neural risk, allowing interventions to target neurodevelopmental trajectories while they are most mutable, and before infant development is substantially impacted [4, 5].

Social cognition and behavior rely on higher-order brain regions that communicate through synchronized neuronal activity [11, 12, 13]. Neural pathology in ASD are thought to disrupt the brain’s ability to generate and sustain synchronous neuronal activity, therefore altering how distributed brain regions communicate with one another [6–10]. Postmortem studies report neural pathology that could disrupt large-scale brain activity in ASD, including differences in neuronal and axonal organization [14, 15], myelination [16], and neurotransmitter receptor density [17]. The large-scale oscillations that emerge from coherent neuronal activity can be directly studied using techniques such as EEG, or indirectly studied using fMRI. Both EEG and fMRI studies suggest that long-range functional connectivity is reduced in children and adults with ASD [18-20]. Although the majority of connectivity differences in ASD have been studied post-diagnosis, the neuronal and synaptic building blocks that scaffold the connectome are established much earlier, during very early brain development. Postmortem studies [21, 22] and human neural stem cell models [23] suggest that the initial stages of neuronal maturation and organization are abnormal in ASD. ASD-associated genes have also been shown to converge upon molecular processes that govern neuronal differentiation and synaptic development [24, 25]. The presence of cellular and synaptic differences during very early brain development could disrupt large-scale oscillatory activity, and therefore be directly captured *in vivo* using EEG.

Characterizing early functional connectivity patterns *in vivo* relies on prospective studies of infants with heightened risk of developing ASD. The younger siblings of children with ASD (familial-risk infants) have an ASD recurrence risk of nearly 20% [26] and, as they are identified based on family history, can be studied from birth. MRI studies report early brain changes in familial-risk infants who later develop ASD. At 6 months of age, differences in structural brain development include atypical white matter integrity across distributed long-range tracts [27] and major tracts such as the corpus callosum [28, 29]. In the same sample of infants, fMRI measures of functional connectivity at 6 months are shown to predict later ASD diagnoses [30]. These data suggest that connectivity is atypical during early infancy in ASD. However, as an indirect measure of neuronal activity, fMRI coactivation patterns cannot assay *how* synchronized neural communication mechanisms are altered. Measuring patterns of functional connectivity using high temporal precision EEG will provide a unique mechanistic insight, complementary to MRI techniques, into neural interactions during infancy in ASD.

EEG is particularly well-suited to clinical screening, as it is portable, relatively low cost, and involves a lower testing burden than MRI [31]. While EEG has been used to study early neural differences in ASD, there have been no multivariate studies that characterize cortex-wide functional connectivity patterns during infancy in ASD. Here we take a data-driven approach to address this gap, mapping differences in functional connectivity at 3 months of age that are associated with later ASD symptoms. Multivariate pattern analysis leverages the rich information provided by neural time series data and represents a powerful way to uncover brain-wide patterns of functional disruptions that lead to ASD. Functional connectivity is quantified through the phase coherence of alpha oscillations (6-12Hz), as alpha coherence is highly sensitive to early neural changes that occur in the context of both typical [32] and atypical brain development [33]. Furthermore, alpha oscillations are specifically associated with the structural [34] and functional [35] properties of long-range connections and may therefore capture earlier markers of the long-range connectivity differences described in children and adults with ASD. Based on previous findings implicating structural differences throughout the corpus callosum during infancy in ASD [28, 29], we hypothesized that atypical interhemispheric coherence would predict a higher level of ASD symptomatology at 18 months.

## Methods and Materials

### Sample

Participants in the present analyses were part of a larger ongoing study examining the development of infants with and without familial risk for ASD across the first 3 years of life. Familial-risk infants had at least one older sibling with a confirmed ASD diagnosis. Initial parent reports of sibling diagnoses were confirmed by a review of documented evidence. Low-risk infants had no reported family history of ASD or other neurodevelopmental disorders within first degree relatives. Infants were recruited from the community through the UCLA Center for Autism Research and Treatment (CART). Sixty-five infants completed an EEG recording session at 3 months and underwent behavioral assessment at 18 months. Demographic details for the sample are provided in Table 1. The study received ethical approval from the relevant institutional review board, and parents provided informed written consent on behalf of all infants in accordance with the Declaration of Helsinki.

**Table 1.** Demographic participant details. While group comparisons were not carried out as part of the present study, sample descriptions are provided according to both familial risk status groupings, and for infants who did (ASD+), and did not (ASD-), meet the cut-off for clinical symptoms on the ADOS-T at 18 months (ADOS-T ≥ 10).

|  | **Familial Risk**<br>(N=36) | **Low Risk**<br>(N=29) | *p* | **ASD+**<br>(N=14) | **ASD-**<br>(N=51) | *p* |
| --- | --- | --- | --- | --- | --- | --- |
| **Sex**<br>n female (% female) | 13 f<br>(36.1%) | 11 f<br>(37.9%) | .54 | 2<br>(14.3%) | 22<br>(43.1%) | .060 |
| **Precise 3 Month EEG age**<br>Mean (SD), Range | 3.18 (0.35),<br>2.57-3.90 | 3.17 (0.32),<br>2.63-4.13 | .87 | 3.05 (.23),<br>2.57-3.43 | 3.21 (.36),<br>2.63-4.13 | .117 |
| **18 Month VDQ**<br>Mean (SD), Range | 43.39 (9.61),<br>21-63.5 | 49.07<br>(10.63),<br>25-66 | .030* | 31.75 (5.82),<br>21-40.50 | 49.95 (7.43),<br>35.5-66 | <.001* |
| **18 Month NVDQ**<br>Mean (SD), Range | 46.07 (7.51),<br>24.5-64 | 51.54 (9.08),<br>25-66 | .011* | 40.96 (9.70),<br>24.5-59.5 | 50.65 (7.02),<br>34-66 | <.001* |
| **18 Month ADOS-T Total Score**<br>Mean (SD), Range | 7.17 (5.12),<br>1-18 | 5.21 (4.44),<br>0-18 | .11 | 14.28 (2.67),<br>10-18 | 4.09 (2.46),<br>0-9 | <.001* |

### ASD Assessment

A trained clinician administered the Toddler Module of the Autism Diagnostic Observation Schedule-Second Edition (ADOS-T) at 18 months of age [36, 37]. The ADOS-T is a gold-standard tool used by clinicians and researchers to assess social-communication and repetitive behaviors in children under 30 months of age. ASD symptoms were quantified using dimensional ADOS-T algorithm scores (total score ranging from 0-18). ADOS-T scores ≥ 10 are highly indicative of a clinically relevant level of symptoms at 18 months and of ASD symptoms at later ages (measured using the ADOS) [38].

### Cognitive Assessment

A trained clinician administered the Mullen Scales of Early Learning (MSEL) [39] at 18 months of age. The MSEL is a standardized measure of developmental abilities for children ranging in age from birth to 68 months of age. Verbal development (VDQ) was calculated from averaged receptive language and expressive language subscale t-scores, and non-verbal development (NVDQ) from averaged visual reception and fine motor subscale t-scores [40].

### EEG Acquisition

Four electrodes close to the eyes (positioned to record electrooculogram (EOG)) were removed from the net to increase comfort for infants. Net Station 4.4.5 software was used to record from a Net Amps 300 amplifier with a low-pass analog filter cutoff frequency of 6 KHz. EEG data were acquired for at least 3 minutes. Data were sampled at 500 Hz and referenced to vertex (Cz) at the time of recording. Electrode impedances were kept below 100 KΩ. Infants were held in a caregiver’s lap throughout the recording while bubbles were blown by an unseen experimenter, consistent with widely used spontaneous recording conditions in infant populations [41, 42].

### EEG Processing

All offline data processing and analyses were performed using EEGLAB [43] and in-house MATLAB scripts. The experimenter was blind to participant details (including risk status) throughout the data cleaning process. Data were high pass filtered to remove frequencies below 1 Hz and low pass filtered to remove frequencies above 90 Hz, using a finite impulse response filter. Continuous data were then visually inspected, and any sections including excessive electromyogram or other non-stereotyped artifacts were removed. Artifact subspace reconstruction (ASR), a data cleaning method that uses sliding window principal component analysis, was then used to remove high amplitude artifacts relative to artifact-free reference data [44]. ASR is especially useful for retaining maximum data in infants (where the length of EEG recordings is limited), as it allows artifacts to be removed while retaining the co-occurring EEG data that represents neural activity. The eeglab function *clean_RawData* was used to implement ASR, with default parameters and rejection threshold *k*=8 [44].

Following interpolation to the international 10–20 system 25 channel montage [45], independent component analysis (ICA) was used to decompose data into maximally independent components (IC), and the power spectral distribution (PSD), scalp topography and time course of each IC were visually examined. IC’s that represented non-neural activity (including EMG, EOG, heart artifact and line noise) were removed from the data.

### Alpha Phase Coherence

Cleaned data were transformed to current source density (CSD) estimates, in order to mitigate the effects of volume conduction [46]. Spherical spline Laplacian transforms were conducted using realistic head geometry, with head radius set at 7cm (representing the average head radius of 3-month-old infants), and flexibility constant m = 3. CSD data were separated into 3-second epochs to obtain coherence metrics. To retain consistent data length across all participants, the first 75 seconds of data were used in all further analyses (representing the minimum data length available across the sample). The newcrossf function provided by eeglab [43] was used to compute phase coherence (ERPCOH) from the aforementioned resting state epochs (for each frequency bin):

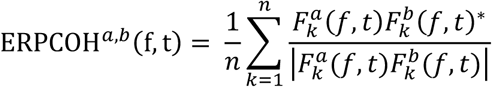

where 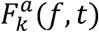 represents the spectral estimate of channel *a* in epoch *k* at frequency *f* and time 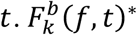 is the complex conjugate of 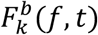 [43]. For each channel pair, ERPCOH was averaged across all frequency bins encompassed by the alpha band (6-12Hz), resulting in 300 values that represented alpha phase coherence between every possible electrode pair.

### Model Fitting

Prediction models were used to assess the relationship between 3-month coherence and 18-month ASD symptoms (ADOS-T total score), with all 300 alpha phase coherence values serving as the initial feature set. A “nested” leave-one-out cross validation (LOOCV) procedure was used whereby one participant was left out of both training and testing samples entirely. A LOOCV regularized regression approach with an elastic net penalty was used to select a subset of functional connections within each fold. Selecting features for each fold while the data of the test subject remained entirely unseen ensured that feature model performance was not falsely inflated through circularity bias [47].

Elastic net regularization is a hybrid approach combining both the ℓ_1_ penalty of lasso and the ℓ_2_ penalty of ridge regression [48, 49], and it is well suited to removing redundant variables and preventing model overfitting for high dimensional data [50]. There are two parameters that impact penalized regression, α and λ. α regulates the degree of mixing between ℓ_1_ and ℓ_2_ penalties, effectively determining the compromise between lasso (least absolute shrinkage and selection operator) and ridge regression techniques. Here we implemented α=0.5 to represent an equal balance between ℓ_1_ and ℓ_2_ penalties. λ is the penalty term and defines the strength of regularization. A geometric sequence of λ values were trialed to determine the λ value that minimized model deviance (mean squared error; MSE), with the final values across all folds averaged to provide a consistent value (λ=1). The *lasso* function in MATLAB was used to implement the regression procedure, and all predictor variables were centered and standardized.

After conducting feature selection within each inner fold, predictor data were centered and scaled, and linear-kernel support vector regression (SVR) models were trained using the default parameters of the *fitrsvm* function in MATLAB. In addition to the advantages of binary classification offered by traditional SVM, support vector machines for regression (SVR) offer an opportunity to assess the value of functional connections for predicting ASD behaviors dimensionally [51]. The resulting model was used to estimate the ADOS-T score of the N=1 participant who defined the validation sample. The procedure was then repeated N times, so that the symptom score for every participant was predicted from a model to which they had not contributed.

Predictive capabilities were examined through the relationship between observed and predicted ADOS-T score. The statistical significance of all LOOCV results was determined using a permutation testing approach [52, 53]. The null distribution of R^2^ was estimated by repeating the entire model fitting procedure (including feature selection within each fold) using 1000 surrogate datasets that were generated under the null hypothesis that there is no relation between 3-month EEG and 18-month ADOS. The final statistical significance of the model was determined by calculating the percentage of null-models that yielded symptom estimates better than the final model. The reported permutation p values therefore represent the probability of observing the reported R^2^ values by chance.

### Predictive Functional Features

A major benefit of multivariate pattern analysis is the ability to examine features that drive the predictive capability of the SVR algorithm. We analyzed the final consensus feature set that consisted of 22 functional connections that had non-zero coefficients in 100% of folds [52], extracting the weight value assigned to each feature. Interpreting the weights from linear models in terms of neural activity patterns can be misleading [54, 55]. To allow neurophysiological interpretation of individual features in the model, SVR weights were transformed into activation patterns using the method described by Haufe and colleagues [55]. Specifically, the activations are derived by,

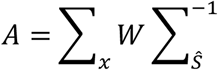

where Σ_x_ denotes the covariance of the data, *W* represents the regression weights, and Σ_s_^−1^ is the inverse covariance of the latent factor.

## Results

### Model Performance

Alpha phase coherence at 3 months predicted ADOS-T scores. Specifically, the SVR model estimated ADOS-T total scores that significantly correlated with actual ADOS-T scores measured at 18 months (*r* = 0.76; *R*^*2*^ = 0.58; *p* = 0.02; see Figure 1). Reported significance values were corrected to represent permutation testing (described under methods).

**Figure 1.**
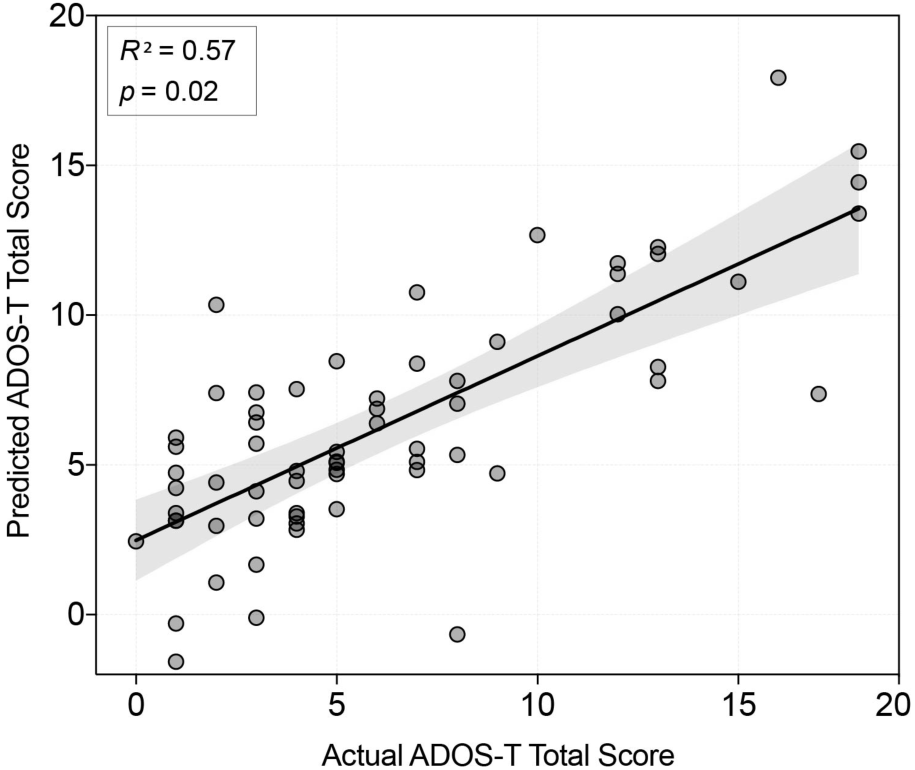
Correlation between actual ADOS-score (X axis), and the predicted ADOS score (Y axis) for each participant, with 95% confidence intervals.

To determine its specificity, we assessed the ability of the model to estimate cognitive function. Consistent with the large overlap between measures of VDQ and ASD symptoms at younger ages [56], ADOS-T scores were more highly correlated with VDQ (*r* = −0.74, *p* <.001) than NVDQ in the present sample (*r* = −0.40, *p* = <.001). We therefore focused on the prediction of NVDQ. Trained on the same input features, the SVR model was unable to predict 18-month NVDQ scores (*r* = 0.15; *p* =0.36). Prediction errors did not vary according to familial-risk group (*p* = 0.20) or sex (*p* = 0.16).

### Feature Activations

As described above, the contribution of individual functional connections to the SVR model was quantified using activation patterns. Activation patterns represent transformed SVR weights that allow neurophysiological interpretation, but do not represent activation patterns as conventionally described in MRI work. Functional connections that contributed to the SVR model represented a mix of positive and negative features (See Figure 2 & 3).

**Figure 2.**
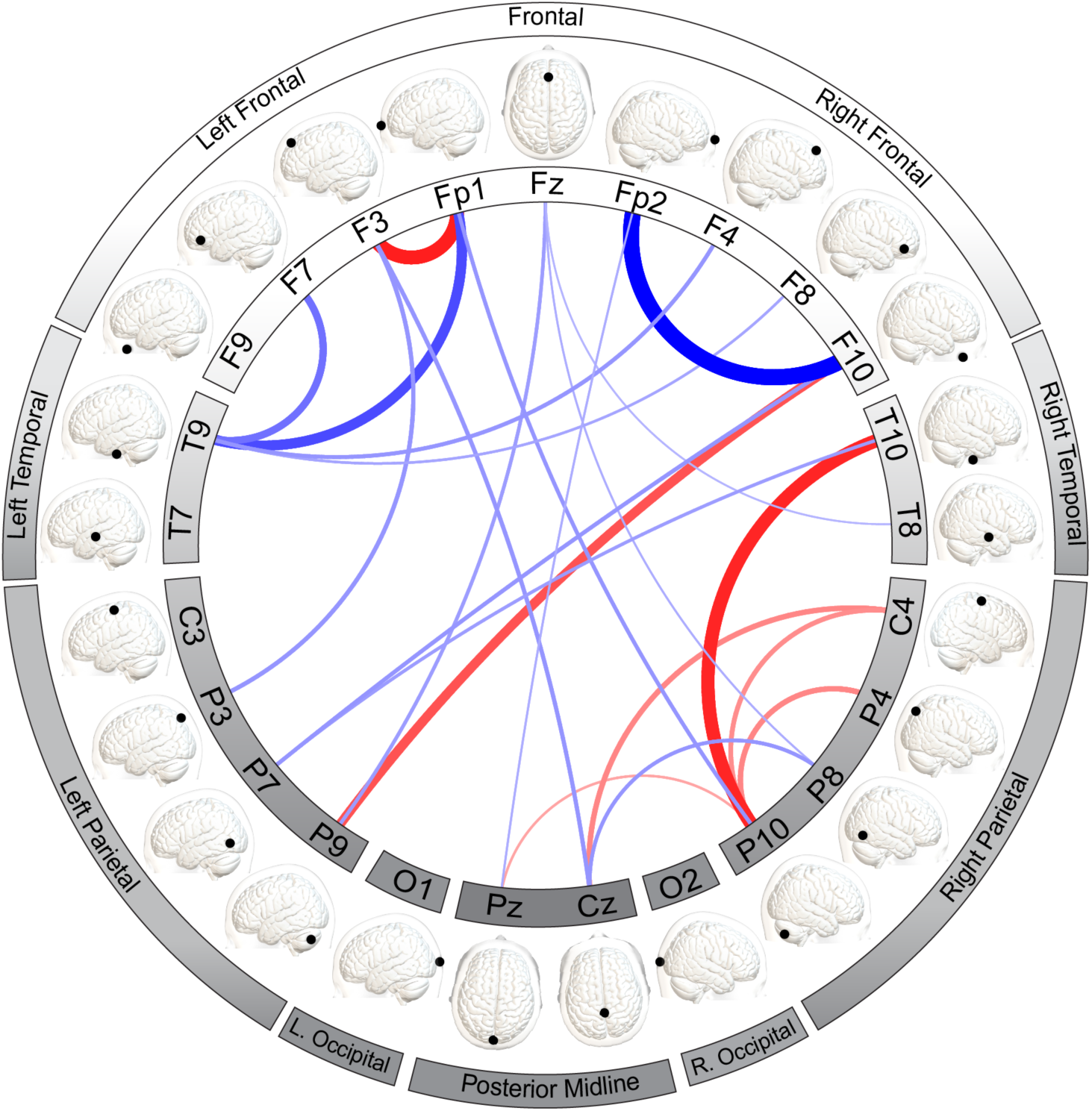
Mean feature activations for each of the 22 predictive function connections that defined the consensus feature set. Red lines represent a positive activation value (higher alpha phase coherence = higher ADOS-T score), and blue lines represent a negative activation value (lower alpha phase coherence = higher ADOS-T score). Wider lines indicating a larger contribution to the model (greater absolute activation strength). Graphical representations indicate the location of each measurement channel.

**Figure 3.**
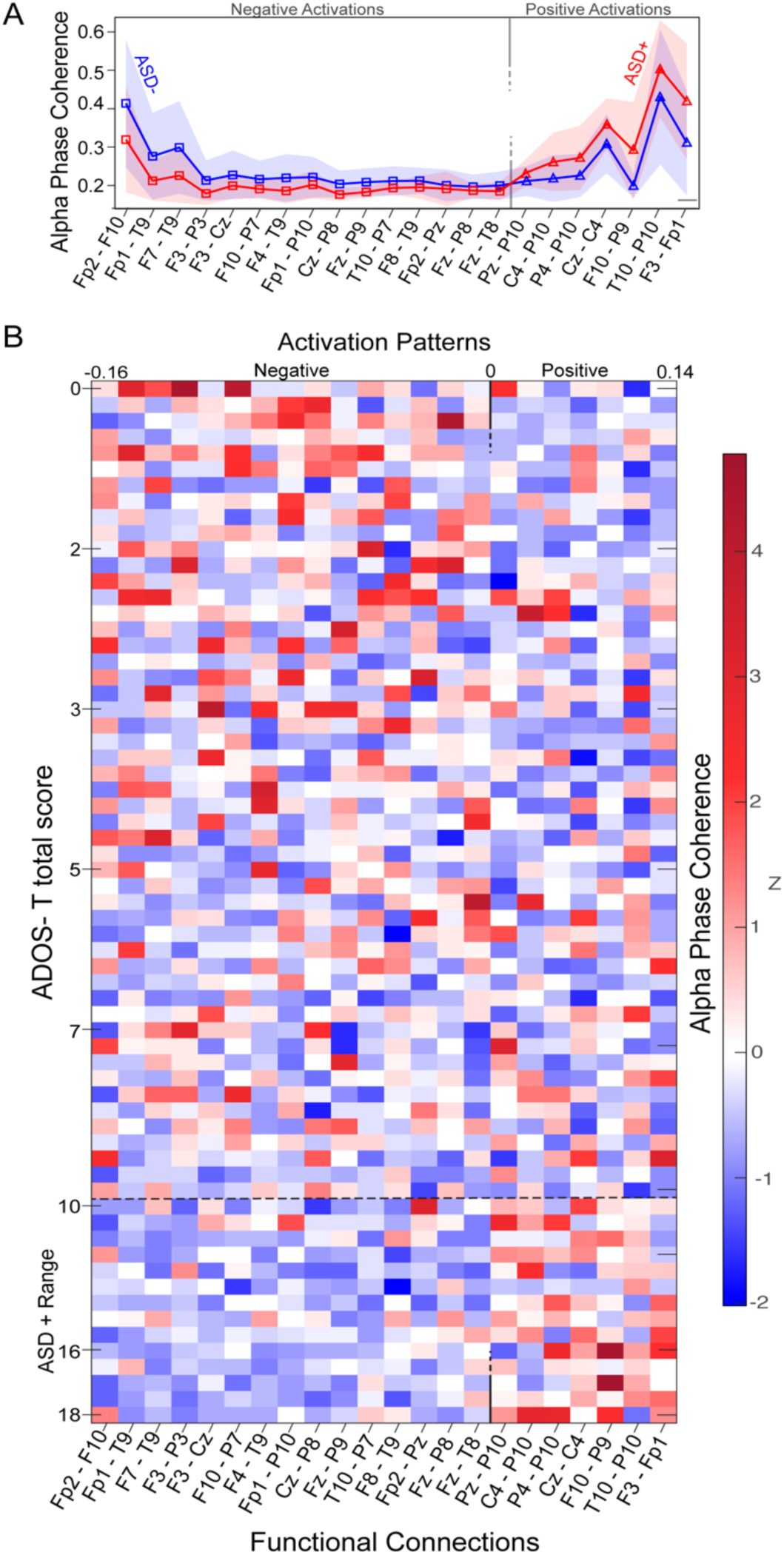
(A) Mean alpha phase coherence for each of the 22 predictive function connections that defined the consensus feature set for ASD+ (red) and ASD- (blue) groups. Shaded regions represent SD. (B) Individual alpha phase coherence values (z scores) for each participant (arranged from low to high ADOS score) for each predictive function connection (with activation patterns arranged from negative to positive).

## Discussion

The present study characterizes functional connectivity patterns during early infancy that predict individual differences in later ASD symptoms. Early connectivity differences that predicted ASD were multivariate, highlighting the importance of studying patterns rather than specific functional connections. The regional distribution of predictive connections shows that decreased connectivity across frontal connections and increased connectivity across temporo-parietal areas are associated with a higher level of ASD symptoms at 18 months. Notably, due to the limited spatial resolution of EEG, the precise cerebral structures driving these results cannot be determined. However, guided by an infant EEG-MRI localization study, we can consider general structures that underlie electrode locations [57].

### Decreased frontal alpha phase coherence

Decreased alpha phase coherence across fronto-frontal, fronto-temporal and fronto-parietal connections predicted higher ASD symptoms. Early disruptions in frontal connectivity are particularly relevant, given the extensive previous literature that implicates frontal neuropathology in ASD. At a cellular level, postmortem studies show disruptions in neuronal [15, 21, 58], axonal [16], laminar [22], and minicolumn [59, 60] organization in the frontal cortex of individuals with ASD. Differences in large scale frontal connectivity (often fronto-posterior hypoconnectivity) are also highly supported by EEG and fMRI studies of children and adults with ASD [20, 61]. We extend these findings to show that frontal disruptions occur prior to behavioral symptoms, suggesting that they represent the core pathophysiology of the disorder, and not simply a consequence of ASD symptoms.

The frontal cortex may be particularly vulnerable to connectivity disruptions in ASD for several reasons, especially given its protracted development [62]. For instance, ASD-associated risk genes are shown to converge upon co-expression networks in the frontal cortex during fetal brain development [24]. By disrupting key neurobiological processes (such as neuronal migration, synaptogenesis, and myelination) in the frontal cortex, genes associated with ASD risk may particularly impact frontal functional connectivity [63]. Further evidence linking these genes to specific frontal disruptions comes from copy number variations and single gene disorders that confer susceptibility for ASD and are also associated with decreased fronto-temporal and fronto-parietal connectivity [64]. The present data suggest that, in addition to the changes seen in syndromic ASD [64], early frontal dysconnectivity due to familial risk may also predispose infants to the emergence of ASD.

### Increased temporo-parietal alpha phase coherence

Positively weighted predictors mainly bridged temporal and parietal areas in the right hemisphere, above brain structures that subserve social information processing [13]: the superior temporal sulcus, as well as postcentral, supramarginal, temporal and angular gyri [57]. These results implicate the right temporoparietal junction (rTPJ) [65], a social hub that coordinates social information processing [66] and shows atypical function in ASD [67]. Alpha phase coherence differences in these regions may reflect the network inefficiencies [27] and structural differences in temporal and parietal white matter tracts that have been identified at 6 months of age in ASD [28], especially considering that white matter integrity is associated with alpha phase coherence [68].

In addition to revealing early connectivity differences during infancy, increased alpha phase coherence in temporal parietal areas may shed mechanistic insight into reports of hypoconnectivity following infancy in ASD. The deleterious effects of increased regional connectivity are well-described in neurocognitive disorders, where periods of increased connectivity are shown to precede decreased connectivity, a pathological process described as hub-overload [69]. Increased alpha phase coherence may lead to hub-overload in ASD and could underlie the transition from over-to under-connectivity seen in both alpha phase coherence and white matter integrity beginning around 2 years of age in ASD [28, 41], as well as widely described reductions in rTPJ activation and connectivity [70].

### Scalability

EEG measures of neural function could serve as scalable and clinically actionable predictors of ASD in early infancy at a time when behavioral signs of atypical development remain unclear. The portability, relatively low cost and low testing burden of EEG renders it practical for community screening in large populations [31]. To translate laboratory-based EEG studies to community settings, neural markers need to be measured accurately under task-free conditions in less controlled environments. Alpha phase coherence, in particular, represents a highly scalable metric. Alpha oscillations are dominant in spontaneous brain activity and are less susceptible to biological and environmental artifacts, thus facilitating measurement in larger clinical or community samples.

### Early Identification & Intervention

Behavioral features that can consistently predict later ASD diagnosis have not been identified in the first year of life and predominantly emerge after 12 months of age [71]. Although EEG is not intended to replace behavioral assessment of ASD, EEG markers are uniquely positioned to elucidate individual differences that confer neural risk for ASD. By examining dimensional risk (rather than binary diagnostic labels), the present study highlights that early network disruptions in ASD occur along a continuum. This approach will facilitate the identification of neural risk associated with milder/borderline ASD symptoms, a clinical group that eludes early behavioral identification [3] but may be particularly responsive to prompt intervention [72].

Early disruptions in brain activity may also impact how an infant responds to their environment, causing a cascading brain-behavior-environment interaction that further impacts brain development [73]. Identifying individuals using objective EEG markers would facilitate a shift from reactionary interventions that focus on modifying established behaviors towards preemptive interventions that may mitigate the effects of early disruptions [74].

### Strengths, Limitations & Future Directions

The present study leveraged the benefits of machine learning to model multivariate data. However, in order to retain interpretable links between neurobiology and behavior, we employed a hypothesis-driven modelling approach that reflects our prioritization of interpretability over prediction. For instance, although the inclusion of additional EEG features may capture interactions leading to better model prediction, focusing on one neurobiologically- and clinically relevant EEG metric (alpha phase coherence) retained our ability to map predictive model features back onto EEG data. These links were also preserved through the use of linear modelling as well as forward modelling transformations [55]. These steps allow us to understand very early brain differences that precede ASD, and ultimately optimize the translatable clinical utility of machine learning methods in ASD.

As with many prior EEG studies of familial-risk infants, a relatively small sample size and lack of independent validation limits the generalizability of this study. To determine if alpha phase coherence patterns can provide a clinically applicable biological marker of risk, we need studies in diverse participant samples representing wider etiological factors beyond familial risk, such as infants with known genetic syndromes or preterm infants as well as a community screened cohort. Another limitation lies in the focus on only one measurement technique. EEG and fMRI provide complementary information about brain function and should be integrated in the future to examine how the timing of structural and functional brain changes relate to one another during the first year of life in ASD. Finally, longitudinal monitoring of behavior, environment, and brain development will broaden our understanding of the dynamic changes in early development in ASD and inform decisions around the exact timing and targets of preventative interventions to ultimately improve developmental outcomes.

## Acknowledgements

The authors wish to thank all of the infants and families who generously participated in this study. The authors are also grateful to Rujuta B. Wilson for her helpful comments on the manuscript.

